# MPRAVarDB: an online database and web server for exploring regulatory effects of genetic variants

**DOI:** 10.1101/2024.04.02.587790

**Authors:** Javlon Nizomov, Weijia Jin, Yi Xia, Yunlong Liu, Zhigang Li, Li Chen

**Author notes:** The authors are equally contributed (Javlon Nizomov, Weijia Jin).

## Abstract

**Summary:** Massively parallel reporter assay (MPRA) is an important technology to evaluate the impact of genetic variants on gene regulation. Here, we present MPRAVarDB, an online database and web server, for exploring regulatory effects of genetic variants. MPRAVarDB harbors 18 MPRA experiments designed to assess the regulatory effects of genetic variants associated with GWAS loci, eQTLs and various genomic features, resulting in a total of 242,818 variants tested across more than 30 cell lines and 30 human diseases or traits. MPRAVarDB empowers the query of MPRA variants by genomic region, disease and cell line or by any combination of these query terms. Notably, MPRAVarDB offers a suite of pretrained machine learning models tailored to the specific disease and cell line, facilitating the genome-wide prediction of regulatory variants. MPRAVarDB is friendly to use, and users only need a few clicks to receive query and prediction results.

**Availability:** https://mpravardb.rc.ufl.edu

**Supplementary information:** Supplementary data are available at *Bioinformatics* online.

## 1 Introduction

Genome wide association studies (GWAS) and expression quantitative trait loci (eQTL) analysis have successfully identified thousands of loci associated with human diseases and traits, most of which are located within the noncoding region. However, within the loci, lead variants are not necessarily the causal variants, while those in tight linkage disequilibrium (LD) may be the causal ones. To address this challenge, massively parallel reporter assays (MPRAs) have been developed, which can perform high-throughput functional screen on tens of thousands of all genetic variants in tight LD with GWAS SNPs or eQTLs to evaluate their impact on gene expression. Since its development, MPRAs have been widely used to test the regulatory effects of genetic variants in tight LD with GWAS SNPs, which are associated with neurodegenerative disease (Cooper *et al*., 2022), immune diseases (Mouri *et al*., 2022), various cancer types such as melanoma and multiple myeloma (Long *et al*., 2022; Ajore *et al*., 2022). In addition, MPRAs have been performed to evaluate the regulatory effects of genetic variants in tight LD with eQTLs identified in GM12878 from the CEU cohort (Abell *et al*., 2022; Tewhey *et al*., 2016) and rare variants from multiple tissues in GTEx (Ferraro *et al*., 2020). Particularly, MPRAs can experimentally test the allelic difference in regulatory effects between alleles (e.g. protective and risk) of the targeted variants in a single experiment. Variants showing significant allelic regulatory activity can be deemed as causal variants (Tewhey *et al*., 2016). Together with the associated risk genes, the identification of these causal variants is a fundamental initial step towards developing a mechanistic understanding that may lead to therapeutic development.

Due to the rapid development of MPRAs, the volume of MPRA data has been growing to a large scale. However, there lacks a centralized repository that can curate data, integrate it, and provide an interactive platform to freely query, retrieve, download and even perform in-depth analysis on it. Existing data repositories such as The NCBI Gene Expression Omnibus (GEO) do not provide tools for interactively exploring the MPRA data, offering only functionality to directly deposit it instead. At the time of writing this manuscript, a web interface named MPRAbase (Zhao *et al*., 2023) is limited to providing the GEO accession number and PubMed ID of the MPRA studies and offering the saturation mutagenesis from promoter and enhancers of ∼20 genes reported by one MPRA study (Kircher *et al*., 2019). Nevertheless, MPRAbase lacks a systematical collection of the MPRA data from a wide range of MPRA studies and lacks an in-detailed curation of MPRA data on the variant level. It is also incapable of offering the facilities to browse, retrieve and download MPRA variants, not to say go beyond the ability of existing MPRA data and perform a genome-wide functional screen for the *de novo* findings. To fill this gap, we introduce MPRAVarDB, an online database and web server, to meet the pressing need for integrating and exploring experimentally validated genetic variants from MPRA studies in public domain. MPRAVarDB harbors 18 MPRA experiments for evaluating the regulatory effects of variants in GWAS loci, eQTLs and genomic features (e.g., 5’UTR, 3’UTR), encompassing a total of 242,818 variants tested across more than 30 cell lines and 30 human diseases or traits. MPRAVarDB empowers the query of MPRA variants by genomic region, disease and cell line of interest or by any combination of these query terms. Importantly, MPRAVarDB provides a set of pretrained machine learning models in the specific context of diseases and cell line, facilitating the genome-wide prediction of regulatory variants. MPRAVarDB is friendly to use, and users only need a few clicks to receive query and prediction results. To the best of our knowledge, MPRAVarDB is the first and most comprehensive web tool developed for this purpose, which will benefit the genetics research community to a great extent.

## 2 Methods

MPRAVarDB consists of two core modules: the database module and the analysis module. The database module collects the processed MPRA summary data from publicly available MPRA experiments, which enables the users to query and download the MPRA variants of interest. The analysis module focuses on providing genome-wide prediction of the regulatory effects for query variants of interest. The backend pretrained machine learning models are developed using the variants curated in the database module. The overview of MPRAVarDB is shown in Fig.1.

### 2.1 MPRA summary data

At the time of writing, MPRAVarDB contains a total of a total of 242,818 variants related to 11,344 nearby genes across 18 MPRA experiments from more than 30 cell lines and 30 human diseases or traits (Supplementary Table 1). The MPRA experiments mainly targets SNPs in tight LD with the GWAS SNPs associated with different diseases or traits and SNPs in tight LD with eQTLs. Each MPRA summary data has been retrieved from the original publications and processed in a unified framework. Details of data collection and processing can be found in the Supplementary Notes. Consequently, three categorical annotations are provided for each variant,

- Variant annotation: genomic coordinate; rsID; reference and alternative allele; genome assembly; description of MPRA variants; description of MPRA study.
- Statistical measurement: log2 fold change, pvalue and FDR of allelic regulatory effect.
- Gene annotation: genomic coordinate of nearest gene; gene symbol; gene strand; distance between variant and nearest gene.

### 2.2 Pretrained machine learning models

We provide three sets of pretrained machine learning models from more than 30 cell lines and 30 diseases or traits, enabling the evaluation of allelic regulatory effects of arbitrary genetic variants that may not exist in the MPRA experiments. The training set for the pretrained model utilizes the processed MPRA summary data based on the selection of MPRA study, cell line and disease. For MPRA studies involving multiple diseases and/or cell lines, we stratify the training data by diseases and/or cell lines and train independent models. The positive set includes MPRA variants with an FDR below 0.1, whereas the negative set consists of MPRA variants with an FDR exceeding 0.8. Specifically, two versions of random forest are provided, which differs in strategies for extracting features from the DNA sequence. The first version of random forest takes the 3-mer frequency of the DNA sequence, which results in 64 features. The second version of random forest utilizes the motif frequency of 633 human motifs (Supplementary Notes). In addition, convolutional neural network (CNN), which employs one-hot encoding DNA sequence, serves as another option. The random forest model is trained using R package “randomForest”, while the CNN is trained using PyTorch 2.2.

### 2.3 Implementation

MPRAVarDB is written in R Shinny. The R Shinny interface is developed and host on a PubApp instance from HiPerGator supercomputer at University of Florida. The PubApp instance consists of 2 CPU cores from AMD EPYC 75F3 Milan 3.0, with each core with 8GM RAM.

### 2.4 Workflow for database module

- **Data preparation:** There are two options for exploring the database. For the “Single Query” option, MPRA study, disease and cell line can be selected from the dropdown list of “Choose a study”, “Choose a disease” and “Choose a cell line”. The query region can be entered into three text boxes named “Chromosome”, “Start Position” and “End Position”. For the “Batch Query” option, a bed file can be uploaded, which contains multiple rows with each row corresponding to a genomic region. Each genomic region consists of three required fields “chromosome”, “start” and “end” and two optional columns “disease” and “cell line”.
- **Data upload:** For the “Batch Query” option, the bed file can be upload by the “Browse” and “Load File” two buttons. A preview option is provided to check the validity of the file format.
- **Select disease and cell line:** For the “Single Query” option, the MPRA study, disease and cell line can be selected from the dropdown list. For the “Batch Query” option, the disease and cell line can be added as two optional columns in the bed file.
- **Results display and download:** For the “Single query” option, once selections are made and the “Run Query” button is clicked, the results will be displayed below in a table. For the “Batch Query” option, once the bed file has been uploaded and the “Run Query” button is clicked, the results will be displayed below in a table. For both options, once the results are displayed, a “Download” button will show up for allowing the users to download the query results.

### 2.5 Workflow for analysis module

- **Model preparation:** The pretrained model is selected based on the choice of reference genome, MPRA study, disease, cell line and types of machine learning models, which can be selected from the dropdown list of “Choose a genome”, “Choose a study” and “Choose a category for stratifying the training data”, “Choose a disease/cell line for model training”, “Choose a model type”, “Choose a feature for prediction (only applied to Random Forest)” and “Choose a file type (BED/FASTA)”.
- **Data preparation:** Two formats of query data are supported: (1) a bed file, which contains multiple variants. Each row corresponds to the genomic coordinate of one query variant, which consists of two required fields “chromosome” and “position”; (2) a fasta file, which includes multiple DNA sequences. Each DNA sequence can be selected surrounding one query variant with a length of 1000bp. Clicking the “Browse” button will upload the query data from the local computer to the online server.
- **Data upload:** The bed or fasta file can be upload by the “Browse” and “Load File” two buttons. A preview option is provided to check the validity of the file format.
- **Prediction, results display and download:** Clicking “Generation Prediction” will call the selected pretrained model to predict the regulatory effects of query variants, which will generate a table with prediction scores. A “Download” button is also provided to download the tables.

## 3 Examples

We present two examples to demonstrate the practical application of the analysis module. The MPRA variant set in the first example is derived from a MPRA study for performing a genome-wide functional screens of 3’UTR variants for human disease and evolution (Griesemer *et al*., 2021), which investigates three sets of variants including disease-associated 3’UTR SNPs in LD from the NHGRI-EBI GWAS catalog, 3’UTR SNPs in regions under positive selection and rare 3’UTR SNPs in genes with outlier expression signatures across tissues in GTEx. Specifically, this set consists of 72,588 3’UTR variants, among which 3549 is considered as the positive set (FDR<0.1) and 43,673 are deemed as the negative set (FDR>0.8). The MPRA variant set in the second example is collected from another MPRA study, which evaluates the disease-associated genetic variants associated with five autoimmune diseases. There are 164 positive variants and 12,664 negative variants in the MPRA variant set. In developing the CNN and random forest models, we randomly sample 80% of positive and negative set and the training set, and treat the rest as the testing set. As a benchmark, we collect the genome-wide precomputed functional score from competing methods, which include CScape (Rogers *et al*., 2017), FATHMM_MKL (Shihab *et al*., 2015), DANN (Quang *et al*., 2015), ReMM (Smedley *et al*., 2016) and FunSeq2 (Fu *et al*., 2014). Consequently, CNN performs best followed by random forest (motif) and random forest (3-mer) (Fig. 1C 1). This observation demonstrates that the advantage of MPRAVarDB’s analysis module, which leverages the disease and cell line-specific MPRA variants in developing a robust machine learning model for genome-wide characterization of functional variants.

**Fig. 1.**
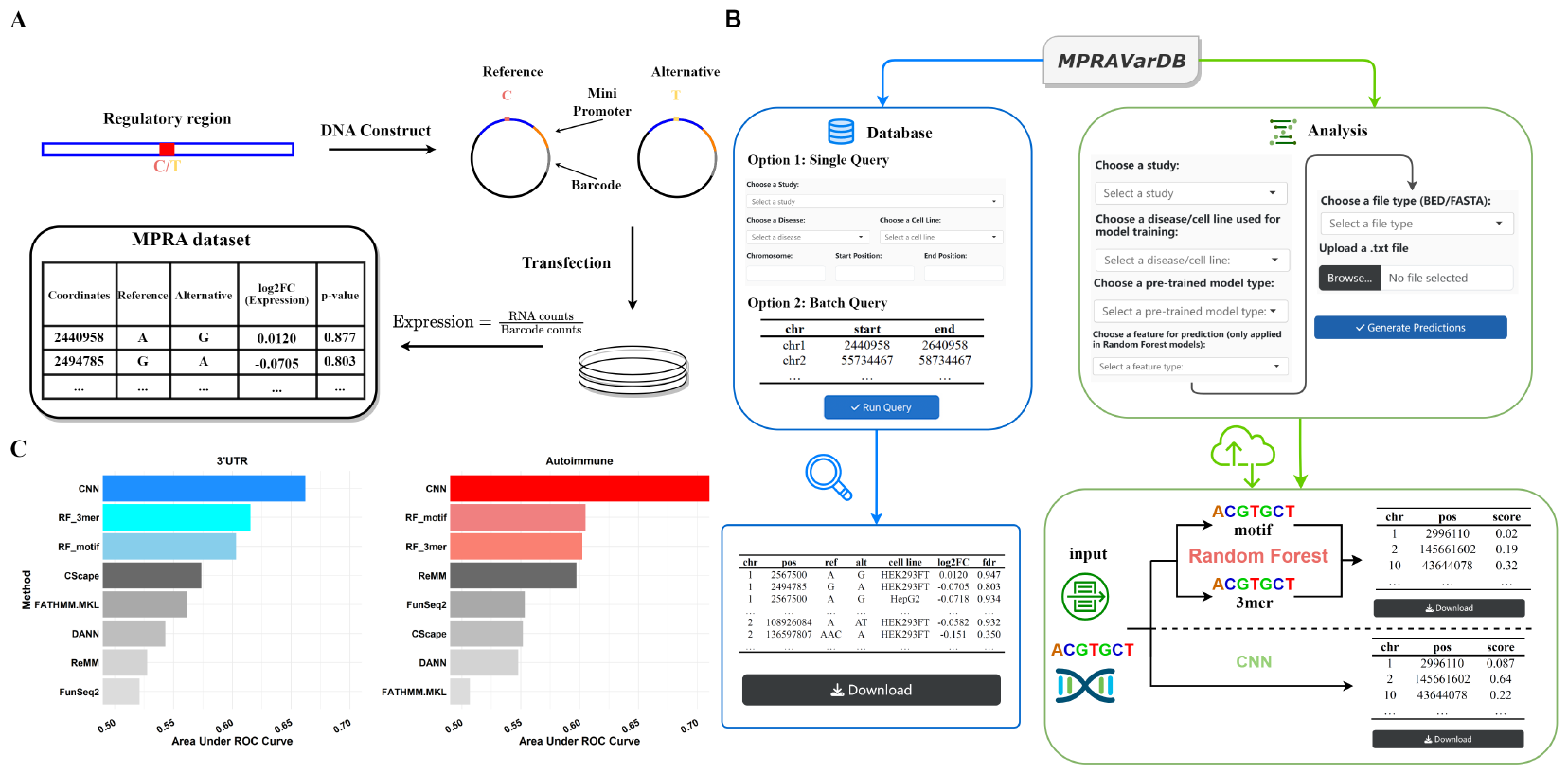
Introduction of the MPRA experiments and overview of MPRAVarDB. (A) A brief introduction of MPRA experiment procedure and summary data. The MPRA experiments start with building the synthetic DNA construct by selecting regulatory sequence that harbor a short polymorphism (e.g., SNP), a minimal promoter and a unique DNA barcode that can be transcribed. Both reference and alternative alleles of the variant are considered in the DNA construct. The surrounding regulatory sequence of the variant may possess the ability to regulate the transcription of the barcode sequence, and each allele may have different ability for the regulation. The DNA constructs are further transfected into cultured cells. After performing RNA and DNA sequencing in the cultured cells, the transcription level is usually estimated as the ratio of RNA counts to barcode counts. To identify variants of significant allelic regulatory effects, statistical analysis is performed to compare the transcriptional activity between the two alleles, which results in a fold change between the transcriptional activity of the two alleles and a binary label, indicating whether the difference is statistically significant (e.g., 1 for FDR<0.1 and 0 otherwise). Variants with significant allelic regulatory effects can be deemed as “causal variants”. (B) Overview of MPRAVarDB. MPRAVarDB consists of the database module and the analysis module. The database module collects the processed MPRA summary data in publicly domains and provide the graphical user interface for users to query and obtain the MPRA variants of interest. The analysis module focus on providing timely prediction for the regulatory effects of query variants by provided pretrained machine learning models, which include random forest and convolutional neural network. Both models are developed based on the labelled variants curated in the database module. (C) Two examples to demonstrate the predictive power of pretrained machine learning models compared to the competing methods.

## 4 Discussion

In this work, we present MPRAVarDB, an online database and web server that enables the exploration of MPRA variants associated with different diseases and cell lines from a wide collection of MPRA studies. The database module of MPRAVarDB facilitates this exploration with a user-friendly interface that allows users to query arbitrary genomic regions. Importantly, the analysis module of MPRAVarDB provides a comprehensive set of pretrained machine learning models, developed on the labelled MPRA variants from the database module, to achieve the prediction for genome-wide genetic variants for evaluating their regulatory efforts. MPRAVarDB reduces a significant obstacle for scientists to retrieve and utilize MPRA variants in public domains. We plan to include more public data as they become available and update the pretrained models accordingly. Our ultimate goal is to turn MPRAVarDB into a powerful toolbox that can efficiently integrate publicly available MPRA variants into a unified framework and make novel prediction on the regulatory effects of variants beyond the experimentally validated ones.

## Acknowledgements

This work was supported by National Institute of General Medical Sciences of the National Institutes of Health under Award Number R35GM142701 to LC.

## Notes

### Competing Interest Statement

The authors have declared no competing interest.

## References

Abell, N. S. et al. (2022). Multiple causal variants underlie genetic associations in humans. Science, 375(6586), 1247–1254.

Ajore, R. et al. (2022). Functional dissection of inherited non-coding variation influencing multiple myeloma risk. Nature communications, 13(1), 151.

Cooper, Y. A. et al. (2022). Functional regulatory variants implicate distinct transcriptional networks in dementia. Science, 377(6608), eabi8654.

Ferraro, N. M. et al. (2020). Transcriptomic signatures across human tissues identify functional rare genetic variation. Science, 369(6509), eaaz5900.

Fu, Y. et al. (2014). Funseq2: a framework for prioritizing noncoding regulatory variants in cancer. Genome Biol, 15(10), 480.

Griesemer, D. et al. (2021). Genome-wide functional screen of 3’utr variants uncovers causal variants for human disease and evolution. Cell, 184(20), 5247–5260.

Kircher, M. et al. (2019). Saturation mutagenesis of twenty disease-associated regulatory elements at single base-pair resolution. Nature communications, 10(1), 3583.

Long, E. et al. (2022). Massively parallel reporter assays and variant scoring identified functional variants and target genes for melanoma loci and highlighted cell-type specificity. The American Journal of Human Genetics, 109(12), 2210–2229.

Mouri, K. et al. (2022). Prioritization of autoimmune disease-associated genetic variants that perturb regulatory element activity in t cells. Nature genetics, 54(5), 603–612.

Quang, D. et al. (2015). Dann: a deep learning approach for annotating the pathogenicity of genetic variants. Bioinformatics, 31(5), 761–3.

Rogers, M. F. et al. (2017). Cscape: a tool for predicting oncogenic single-point mutations in the cancer genome. Sci Rep, 7(1), 11597.

Shihab, H. A. et al. (2015). An integrative approach to predicting the functional effects of non-coding and coding sequence variation. Bioinformatics, 31(10), 1536–1543.

Smedley, D. et al. (2016). A whole-genome analysis framework for effective identification of pathogenic regulatory variants in mendelian disease. The American Journal of Human Genetics, 99(3), 595–606.

Tewhey, R. et al. (2016). Direct identification of hundreds of expression-modulating variants using a multiplexed reporter assay. Cell, 165(6), 1519–1529.

Zhao, J. et al. (2023). Mprabase: A massively parallel reporter assay database. bioRxiv, pages 2023–11.

