## Supplementary materials for "MPRAVarDB: an online database and web server for exploring regulatory effects of genetic variants"

### Supplementary table

#### Table1: Summary of 18 MPRA studies

| Study | disease | Cell line | Description |
|---|---|---|---|
| Functional regulatory variants implicate distinct transcriptional networks in dementia (Cooper et al., 2022) | Alzheimer's disease (AD) and Progressive supranuclear palsy (PSP) | HEK293T | 5706 noncoding single-nucleotide variants, representing 25 genome-wide significant loci associated with AD and nine loci associated with PSP |
| Multiple causal variants underlie genetic associations in humans (Abell et al., 2022) | None | GM12878 | 30,893 variants in LD with selected independent, common, and top- ranked eQTL across 744 eGenes identified in the CEU cohort |
| Transcriptomic signatures across human tissues identify functional rare genetic variation (Ferraro et al., 2020) | None | GM12878 | Rare genetic variants contributing to extreme patterns in gene expression, allelic expression, and alternative splicing across 49 tissues in GTEx |
| Genome-wide functional screen of 3'UTR variants uncovers causal variants for human disease and evolution (Griesemer et al., 2021) | NHGRI-EBI GWAS catalog | HMEC,HEK293FT,HEPG2,K562,GM12878,SKNSH | 3' UTR SNPs and insertion/ deletions (indels) in LD from the NHGRI-EBI GWAS catalog, which include 2,153 putative disease-associated variants including from 1,556 independent association loci; 9,325 3'UTR SNPs and indels overlapping regions identified as being under positive selection in humans; a set of 46 rare 3'UTR variants in genes with outlier expression signatures across tissues in GTEx. |
| Functional testing of thousands of osteoarthritis-associated variants for regulatory activity (Klein et al., 2019) | Osteoarthritis | Saos-2 | 1,605 SNPs in high LD (r2 > 0.8) at 35 lead SNPs associated with OA in Europeans via GWAS |
| Transcriptional-regulatory convergence across functional MDD risk variants identified by massively parallel reporter assays (Mulvey & Dougherty, 2021) | Major depressive disorder (MDD) | N2A | Over 1000 SNPs are selected from 39 neuropsychiatric trait/disease GWAS loci, which is based on overlap with eQTL and histone marks from human brains and cell cultures. |
| Saturation mutagenesis of twenty disease-associated regulatory elements at | ~20 diseases and details can be found at | HepG2 (HB-8065), HEK293T | Over 30,000 single nucleotide substitutions and deletions in 20 disease-associated gene promoters and enhancers |

| | | | |
|---|---|---|---|
| single base-pair resolution (Kircher et al., 2019) | https://kircherlab.bihealth.org/satMutMPRA/ | (CRL-11268), HeLa (CCL-2), HaCaT (CRL-2404), Neuro-2a (CCL-131), LNCaP (CRL-1740), and SK-MEL-28 (HTB-72) | |
| Functional dissection of inherited non-coding variation influencing multiple myeloma risk (Ajore et al., 2022) | Multiple myeloma (MM) | L363, MOLP8 | 1039 variants in high LD (r2 > 0.8) at 23 MM risk loci |
| Global discovery of lupus genetic risk variant allelic enhancer activity (Lu et al., 2021) | Systemic Lupus Erythematosus (SLE) | GM12878, Jurkat T cell | 3073 variants in high LD (r2 > 0.8) at 91 SLE-associated risk loci |
| Prioritization of autoimmune disease-associated genetic variants that perturb regulatory element activity in T cells (Mouri et al., 2022) | Five Autoimmune diseases: Type 1 diabetes (T1D), Ulcerative colitis (UC),Crohn's disease (CD), Rheumatoid arthritis (RA), Multiple sclerosis (MS). | Jurkat T cell | 18,312 variants in tight LD (r2>0.8) with 578 GWAS index variants representing 531 distinct GWAS loci. |
| Massively parallel reporter assays and variant scoring identified functional variants and target genes for melanoma loci and highlighted cell-type specificity (Long et al., 2022) | Melanoma | UACC903 (malignant melanoma) C283T (normal melanocyte cells) | 1,992 risk-associated variants in tight LD (r2>0.8) from 54 risk-associated loci |
| Massively parallel reporter assays of melanoma risk variants identify MX2 as a gene promoting melanoma (Choi et al., 2020) | Melanoma | UACC903, HEK293FT | 832 high-LD variants at melanoma susceptibility GWAS loci |
| A screen of 1049 schizophrenia and 30 Alzheimer's-associated variants for regulatory potential (Myint et al., 2020) | Alzheimer's disease and Schizophrenia | K562,SK-SY5Y | 1,049 SZ and 30 AD variants in 64 SZ loci and 9 AD loci, respectively |

| | | | |
|---|---|---|---|
| Systematic Functional Dissection of Common Genetic Variation Affecting Red Blood Cell Traits (Ulirsch et al., 2016) | Red Blood Cell Traits | K562, K562+GATA1 | 2,756 variants in strong linkage disequilibrium with 75 sentinel variants associated with red blood cell traits |
| Direct Identification of Hundreds of Expression-Modulating Variants using a Multiplexed Reporter Assay (Tewhey et al., 2016) | NA | GM12878 | 32,373 variants associated with eQTLs in lymphoblastoid cell lines |
| Allele-specific expression and high-throughput reporter assay reveal functional genetic variants associated with alcohol use disorders (Rao et al., 2021) | Alcohol use disorders | SH-SY5Y and SK-N-BE(2) | SNPs)] in 3'UTR of 88 genes identified from allele-specific expression analysis between 30 subjects with Alcohol use disorders and 30 controls. |
| Multi-level functional genomics reveals molecular and cellular oncogenicity of patient-based 3' untranslated region mutations(Schuster et al., 2023) | Prostate cancer | PC3 cell | 14,497 single-nucleotide mutations enriched in oncogenic pathways and 30 UTR regulatory elements. |
| Systematic investigation of allelic regulatory activity of schizophrenia-associated common variants (McAfee et al., 2023) | Schizophrenia | HEK293s,HNPS | 5,173 fine-mapped schizophrenia GWAS variants |

### Table2: Summary of 18 annotations for each variant

| Column name | Interpretation |
|---|---|
| chr | chromosome of the variant |
| pos | genomic position of the variant |
| ref | reference allele of the variant |
| alt | alternative allele of the variant |
| genome | reference genome of the variant |
| rsid | SNP ID of the variant |
| disease | disease type from the GWAS |
| cell line | cell line used from MPRA experiment |
| description | brief description of the source of MPRA variants |
| log2FC | fold change of expression change between reference and alternative allele |
| pvalue | pvalue of expression change between reference and alternative allele |
| fdr | fdr of expression change between reference and alternative allele |
| distancetoGene | distance of the variant to the nearest gene |
| insideGene | variant inside the nearest gene or not |
| gene_name | nearest gene name |

| | |
|---|---|
| gene_start | genomic start position of the nearest gene |
| gene_end | genomic end position of the nearest gene |
| gene_strand | strand of the nearest gene |
| MPRA study | name of the MPRA study |

### Supplementary Notes

**Computational pipeline to process MPRA summary data**

We collect the MPRA summary data from 18 MPRA studies in Table 1. Next, we use a computational pipeline to process MRPA variants from all studies into the same format with 18 annotations (Table 2). For annotations missing from the MPRA summary data, we will adopt the following bioinformatics tools to generate the annotations. Specifically,

1. **Variant annotation:**
   If genomic position, reference and alternative alleles are provided for variants in the MPRA summary data but rsID is missing, we adopt "snpsByOverlaps" function in in "SNPlocs.Hsapiens.dbSNP144.GRCh37/GRCh38" R/Bioconductor package, which takes the "GRanges" object of query variants and reference SNPs in "SNPlocs.Hsapiens.dbSNP144.GRCh37/GRCh38" R/Bioconductor package as the parameters and output the rsID. If rsID is provided for variants in the MPRA summary data but reference and alternative alleles are missing, we adopt "snpsById" function, which takes rsID and reference SNPs in "SNPlocs.Hsapiens.dbSNP144.GRCh37/GRCh38" R/Bioconductor package as the parameters and output the genomic positions of query variants. Next, we adopt "inferRefAndAltAlleles" function in "BSgenome.Hsapiens.UCSC.hg19/hg38" R/Bioconductor package, which takes the genomic positions of query variants and human refence genome in "BSgenome.Hsapiens.UCSC.hg19/hg38" R/Bioconductor package as the parameters and output the reference and alternative alleles for the query variants.

2. **Statistical measurement:**
   If log2FC and pvalue are missing, we fill the missing entries with NA.

3. **Gene annotation:**
   Entrez IDs for all human protein-coding genes are retrieved from "TxDb.Hsapiens.UCSC.hg19/hg38.knownGene" R/Bioconductor package. Accordingly, gene symbols are obtained from R/Bioconductor package "org.Hs.eg.db" by using "select" function with Entrez IDs as the parameter. Next, gene annotations can be obtained by using "annotatePeakInBatch" function in R/Bioconductor package "ChIPpeakAnno" with "GRanges" object of query variants and reference gene annotations in "EnsDb.Hsapiens.v75/v86" R/Bioconductor package as the parameters and output the gene-level annotations for query variants.

**Feature extraction for machine learning models**

1. **Motif frequency for DNA sequence**
   For random forest model with motif frequency as the feature, we obtain the motif feature in the R/Bioconductor environment. The human motifs are collected from Bioconductor package "JASPAR2020" with species ID 9606, which results in 633 Position frequency matrix (PFM). Motif scan is carried out by using the 633 PFMs to scan 1kb DNA sequence surrounding each MPRA variant with the variant in the middle. The DNA sequence can be provided in the fasta file directly or as genomic regions in the bed file. If input file is fasta file, the DNA sequences will be converted into "DNAStringSet" object in R/Bioconductor package "Biostrings". If input file is bed file, the genomic regions will be converted to "GRanges" object in R/Bioconductor package "GenomicRanges". Next, "matchMotifs" function in R/Bioconductor package "motifmatchr", if input file is fasta file, takes two parameters including (1) 633 human motifs in the format of PFMs (2) DNA sequences stored in the "DNAStringSet" object; if input file is bed file, takes three parameters including (1) 633 human motifs in the format of PFMs (2) genomic regions stored in a "GRanges" object and (3) Human reference genome represented R/Bioconductor package"BSgenome.Hsapiens.UCSC.hg19/hg38". As a result, for each DNA sequence, the motif features will be generated as a vector of 633 motif frequency.

2. **3mer frequency for DNA sequence**
   For random forest with 3mer frequency as the feature, we obtain 3mer frequency in the R/Bioconductor environment. If input file is fasta file, the DNA sequences will be converted into "DNAStringSet" object in R/Bioconductor package "Biostrings". If the input file is bed file, the genomic regions will be converted to "GRanges" object in R/Bioconductor package "GenomicRanges". "getSeq" function in R/Bioconductor package "BSgenome", takes both the "GRanges" object and human reference genom"BSgenome.Hsapiens.UCSC.hg19/hg38" as the parameters and output the DNA sequences in a "DNAStringSet" object. Next, "trinucleotideFrequency" function in R/Bioconductor package "Biostrings" takes "DNAStringSet" object and extracts the 3mer frequency, resulting in 64 3mer features.

3. **One-hot encoding for DNA sequence**
   For the convolutional neural network (CNN), 1kb DNA sequence surrounding each MPRA variant with the variant in the middle is one-hot encoded (A: [1,0,0,0], C: [0,1,0,0], G: [0,0,1,0] and T: [0,0,0,1]) as the feature representation for each MPRA variant.

### Reference


Abell, N. S., DeGorter, M. K., Gloudemans, M. J., Greenwald, E., Smith, K. S., He, Z., & Montgomery, S. B. (2022). Multiple causal variants underlie genetic associations in humans. *Science*, *375*(6586), 1247-1254. https://doi.org/10.1126/science.abj5117

Ajore, R., Niroula, A., Pertesi, M., Cafaro, C., Thodberg, M., Went, M., Bao, E. L., Duran-Lozano, L., Lopez de Lapuente Portilla, A., Olafsdottir, T., Ugidos-Damboriena, N., Magnusson, O., Samur, M., Lareau, C. A., Halldorsson, G. H., Thorleifsson, G., Norddahl, G. L., Gunnarsdottir, K., Forsti, A., . . . Nilsson, B. (2022). Functional dissection of inherited non-coding variation influencing multiple myeloma risk. *Nat Commun*, *13*(1), 151. https://doi.org/10.1038/s41467-021-27666-x

Choi, J., Zhang, T., Vu, A., Ablain, J., Makowski, M. M., Colli, L. M., Xu, M., Hennessey, R. C., Yin, J., Rothschild, H., Grawe, C., Kovacs, M. A., Funderburk, K. M., Brossard, M., Taylor, J., Pasaniuc, B., Chari, R., Chanock, S. J., Hoggart, C. J., . . . Brown, K. M. (2020). Massively parallel reporter assays of melanoma risk variants identify MX2 as a gene promoting melanoma. *Nat Commun*, *11*(1), 2718. https://doi.org/10.1038/s41467-020-16590-1

Cooper, Y. A., Teyssier, N., Drager, N. M., Guo, Q., Davis, J. E., Sattler, S. M., Yang, Z., Patel, A., Wu, S., Kosuri, S., Coppola, G., Kampmann, M., & Geschwind, D. H. (2022). Functional regulatory variants implicate distinct transcriptional networks in dementia. *Science*, *377*(6608), eabi8654. https://doi.org/10.1126/science.abi8654

Ferraro, N. M., Strober, B. J., Einson, J., Abell, N. S., Aguet, F., Barbeira, A. N., Brandt, M., Bucan, M., Castel, S. E., Davis, J. R., Greenwald, E., Hess, G. T., Hilliard, A. T., Kember, R. L., Kotis, B., Park, Y., Peloso, G., Ramdas, S., Scott, A. J., . . . Battle, A. (2020). Transcriptomic signatures across human tissues identify functional rare genetic variation. *Science*, *369*(6509). https://doi.org/10.1126/science.aaz5900

Griesemer, D., Xue, J. R., Reilly, S. K., Ulirsch, J. C., Kukreja, K., Davis, J. R., Kanai, M., Yang, D. K., Butts, J. C., Guney, M. H., Luban, J., Montgomery, S. B., Finucane, H. K., Novina, C. D., Tewhey, R., & Sabeti, P. C. (2021). Genome-wide functional screen of 3'UTR variants uncovers causal variants for human disease and evolution. *Cell*, *184*(20), 5247-5260 e5219. https://doi.org/10.1016/j.cell.2021.08.025

Kircher, M., Xiong, C., Martin, B., Schubach, M., Inoue, F., Bell, R. J. A., Costello, J. F., Shendure, J., & Ahituv, N. (2019). Saturation mutagenesis of twenty disease-associated regulatory elements at single base-pair resolution. *Nat Commun*, *10*(1), 3583. https://doi.org/10.1038/s41467-019-11526-w

Klein, J. C., Keith, A., Rice, S. J., Shepherd, C., Agarwal, V., Loughlin, J., & Shendure, J. (2019). Functional testing of thousands of osteoarthritis-associated variants for regulatory activity. *Nat Commun*, *10*(1), 2434. https://doi.org/10.1038/s41467-019-10439-y

Long, E., Yin, J., Funderburk, K. M., Xu, M., Feng, J., Kane, A., Zhang, T., Myers, T., Golden, A., Thakur, R., Kong, H., Jessop, L., Kim, E. Y., Jones, K., Chari, R., Machiela, M. J., Yu, K., Melanoma Meta-Analysis, C., Iles, M. M., . . . Choi, J. (2022). Massively parallel reporter assays and variant scoring identified functional variants and target genes for melanoma loci and highlighted cell-type specificity. *Am J Hum Genet*, *109*(12), 2210-2229. https://doi.org/10.1016/j.ajhg.2022.11.006

Lu, X., Chen, X., Forney, C., Donmez, O., Miller, D., Parameswaran, S., Hong, T., Huang, Y., Pujato, M., Cazares, T., Miraldi, E. R., Ray, J. P., de Boer, C. G., Harley, J. B., Weirauch, M. T., & Kottyan, L. C. (2021). Global discovery of lupus genetic risk variant allelic enhancer activity. *Nat Commun*, *12*(1), 1611. https://doi.org/10.1038/s41467-021-21854-5

McAfee, J. C., Lee, S., Lee, J., Bell, J. L., Krupa, O., Davis, J., Insigne, K., Bond, M. L., Zhao, N., Boyle, A. P., Phanstiel, D. H., Love, M. I., Stein, J. L., Ruzicka, W. B., Davila-Velderrain, J., Kosuri, S., & Won, H. (2023). Systematic investigation of allelic regulatory activity of schizophrenia-associated common variants. *Cell Genom*, *3*(10), 100404. https://doi.org/10.1016/j.xgen.2023.100404

Mouri, K., Guo, M. H., de Boer, C. G., Lissner, M. M., Harten, I. A., Newby, G. A., DeBerg, H. A., Platt, W. F., Gentili, M., Liu, D. R., Campbell, D. J., Hacohen, N., Tewhey, R., & Ray, J. P. (2022). Prioritization of autoimmune disease-associated genetic variants that perturb regulatory element activity in T cells. *Nat Genet*, *54*(5), 603-612. https://doi.org/10.1038/s41588-022-01056-5

Mulvey, B., & Dougherty, J. D. (2021). Transcriptional-regulatory convergence across functional MDD risk variants identified by massively parallel reporter assays. *Transl Psychiatry*, *11*(1), 403. https://doi.org/10.1038/s41398-021-01493-6

Myint, L., Wang, R., Boukas, L., Hansen, K. D., Goff, L. A., & Avramopoulos, D. (2020). A screen of 1,049 schizophrenia and 30 Alzheimer's-associated variants for regulatory potential. *Am J Med Genet B Neuropsychiatr Genet*, *183*(1), 61-73. https://doi.org/10.1002/ajmg.b.32761

Rao, X., Thapa, K. S., Chen, A. B., Lin, H., Gao, H., Reiter, J. L., Hargreaves, K. A., Ipe, J., Lai, D., Xuei, X., Wang, Y., Gu, H., Kapoor, M., Farris, S. P., Tischfield, J., Foroud, T., Goate, A. M., Skaar, T. C., Mayfield, R. D., . . . Liu, Y. (2021). Allele-specific expression and high-throughput reporter assay reveal functional genetic variants associated with alcohol use disorders. *Mol Psychiatry*, *26*(4), 1142-1151. https://doi.org/10.1038/s41380-019-0508-z

Schuster, S. L., Arora, S., Wladyka, C. L., Itagi, P., Corey, L., Young, D., Stackhouse, B. L., Kollath, L., Wu, Q. V., Corey, E., True, L. D., Ha, G., Paddison, P. J., & Hsieh, A. C. (2023). Multi-level functional genomics reveals molecular and cellular oncogenicity of patient-based 3' untranslated region mutations. *Cell Rep*, *42*(8), 112840. https://doi.org/10.1016/j.celrep.2023.112840

Tewhey, R., Kotliar, D., Park, D. S., Liu, B., Winnicki, S., Reilly, S. K., Andersen, K. G., Mikkelsen, T. S., Lander, E. S., Schaffner, S. F., & Sabeti, P. C. (2016). Direct Identification of Hundreds of Expression-Modulating Variants using a Multiplexed Reporter Assay. *Cell*, *165*(6), 1519-1529. https://doi.org/10.1016/j.cell.2016.04.027

Ulirsch, J. C., Nandakumar, S. K., Wang, L., Giani, F. C., Zhang, X., Rogov, P., Melnikov, A., McDonel, P., Do, R., Mikkelsen, T. S., & Sankaran, V. G. (2016). Systematic Functional Dissection of Common Genetic Variation Affecting Red Blood Cell Traits. *Cell*, *165*(6), 1530-1545. https://doi.org/10.1016/j.cell.2016.04.048
